# *Dbx2*, an aging-related homeobox gene, inhibits the proliferation of adult neural progenitors

**DOI:** 10.1101/2023.01.16.524218

**Authors:** Andrea Giuliani, Valerio Licursi, Paola S. Nisi, Mario Fiore, Stefano Biagioni, Rodolfo Negri, Peter J. Rugg-Gunn, Emanuele Cacci, Giuseppe Lupo

## Abstract

The subventricular zone (SVZ) of the adult mouse brain contains quiescent neural stem cells, which can be activated (aNSCs) to generate transit amplifying progenitors (TAPs), neuroblasts (NBs) and newborn neurons. Neurogenesis declines during aging, as the aged SVZ niche causes transcriptomic changes that promote NSC quiescence and decrease proliferating neural/stem progenitor cells (NSPCs). The transcription factors mediating these changes, however, remain unclear. We previously found that the homeobox gene *Dbx2* is upregulated in aged SVZ NSPCs and inhibits NSPC culture growth. Here, we report that *Dbx2* is repressed by Epidermal Growth Factor Receptor signaling, which promotes NSPC proliferation and decreases in the aged SVZ. We show that *Dbx2* inhibits NSPC proliferation by hindering the G2/M transition and elucidate the transcriptomic networks modulated by *Dbx2*, highlighting its role in the downregulation of the cell cycle molecular pathways. Accordingly, *Dbx2* function is negatively correlated with the transcriptional signatures of proliferative NSPCs (aNSCs, TAPs and early NBs). These results point to *Dbx2* as a molecular node relaying the anti-neurogenic input of the aged niche to the NSPC transcriptome.

## INTRODUCTION

In adult mice, the subventricular zone (SVZ) of the lateral ventricles and the subgranular zone (SGZ) of the dentate gyrus harbour neurogenic neural stem/progenitor cells (NSPCs) thanks to a specialized niche supporting NSPC maintenance and neuronal differentiation. SVZ and SGZ NSPCs are organized in a cell lineage encompassing the neurogenic process (Kempermann *et al*, 2015)(Lim & Alvarez-Buylla, 2016). Neural stem cells (NSCs) lie at the top of the lineage and are largely quiescent (qNSCs). Upon activation (aNSCs) by appropriate stimuli, they can enter the cell cycle, and self-renew or generate daughter cells that give rise to transient amplifying progenitors (TAPs), neuroblasts (NBs) and newborn neurons (Obernier & Alvarez-Buylla, 2019)(Urbán *et al*, 2019).

Adult neurogenesis drops with age, which may be relevant to the cognitive decline of elderly humans (Moreno-Jiménez *et al*, 2019)(Terreros-Roncal *et al*, 2021). This is best known in mice, wherein proliferating NSPCs and their neuronal output are markedly decreased in the aged SVZ and SGZ (Lupo *et al*, 2019b)(Navarro Negredo *et al*, 2020). Two explanations have been proposed for this reduction. Earlier models suggest that the age-dependent decrease in neurogenesis is driven by the progressive exhaustion of the qNSC pool due to their activation and differentiation (Encinas *et al*, 2011)(Calzolari *et al*, 2015). According to recent studies, however, a qNSC reservoir persists in the aged brain (Bast *et al*, 2018)(Kalamakis *et al*, 2019)(Xie *et al*, 2020)(Ibrayeva *et al*, 2021). Moreover, some NSCs may return to quiescence after activation and self-renewal (Basak *et al*, 2018)(Belenguer *et al*, 2021)(Harris *et al*, 2021). These observations lead to alternative models, whereby the neurogenic decline would be caused by the increased quiescence of the aged NSC pool. Considering the heterogeneous nature of NSC populations (Kalamakis *et al*, 2019)(Xie *et al*, 2020)(Ibrayeva *et al*, 2021), and the time-dependent changes in neurogenesis (Apostolopoulou *et al*, 2017)(Harris *et al*, 2021), both models may be represented in certain NSC subpopulations and/or ages.

Adult NSPC proliferation is influenced by various extracellular signals. In the mouse SVZ, Epidermal Growth Factor receptor (EGFR) signaling is crucial in qNSC activation and is reduced in the aged niche (Enwere *et al*, 2004)(Codega *et al*, 2014)(Kobayashi *et al*, 2019)(Cochard *et al*, 2021). In contrast, inflammatory pathways, such as interferon (IFN) signaling, increase in the aged SVZ and promote NSC quiescence (Dulken *et al*, 2019)(Kalamakis *et al*, 2019). Tumor Necrosis Factor α (TNFα), Transforming Growth Factor β (TGFβ) and Fibroblast Growth Factor (FGF) pathways are also implicated in NSC quiescence/activation and aging (Pineda *et al*, 2013)(Yamada *et al*, 2017) (Belenguer *et al*, 2021)(Marqués-Torrejón *et al*, 2021).

Transcriptomic comparison of qNSCs and aNSCs has revealed clear differences in the expression of genes related to cell cycle, protein synthesis and inflammatory response (Codega *et al*, 2014) (Belenguer *et al*, 2021)(Harris *et al*, 2021). Transcriptomic changes have also been detected between young adult and aged NSPCs (Shi *et al*, 2018)(Leeman *et al*, 2018)(Lupo *et al*, 2018)(Kalamakis *et al*, 2019)(Xie *et al*, 2020)(Ibrayeva *et al*, 2021). This suggests that transcriptional regulation in NSPCs is key to control their proliferation and to the alterations of aged NSPCs, but the transcription factors relaying aged niche signals to the NSPC transcriptome remain unclear.

We previously showed that the homeobox gene *Dbx2* is upregulated in NSPCs of the aged mouse SVZ and inhibits the growth of NSPC cultures (Lupo *et al*, 2018), but the extracellular signals controlling *Dbx2* expression and the mechanisms mediating *Dbx2* function remained unclear. In this work, we show that *Dbx2* transcription is negatively modulated by EGRF signaling. We provide evidence that *Dbx2* can hinder the G2/M transition in adult NSPCs, thus reducing their proliferation. Finally, we describe the *Dbx2*-regulated transcriptome in adult NSPC cultures. Notably, this includes cell cycle and inflammatory response gene networks, and is negatively correlated with the signatures of the proliferative neurogenic cell types (aNSCs, TAPs and early NBs).

## RESULTS AND DISCUSSION

### *Dbx2* expression is upregulated in aged or quiescent NSPCs and is repressed by EGFR signaling

We previously reported that *Dbx2* mRNA levels were increased both in NSPC cultures and in freshly sorted NSPCs obtained from the SVZ of aged (18 mo) mice as compared to young adult (3-7 mo) samples (Lupo *et al*, 2018). To confirm these results, we checked a recent transcriptomic dataset of NSPCs bulk-sorted from the SVZ of 2 mo, 7 mo and 19 mo mice, noticing an age-dependent upregulation of *Dbx2* (Kalamakis *et al*, 2019) (Fig 1A). We also examined published transcriptomic datasets of qNSCs and aNSCs bulk-sorted from the adult mouse SVZ (Codega *et al*, 2014)(Xie *et al*, 2020)(Belenguer *et al*, 2021), consistently observing *Dbx2* upregulation in qNSCs (Fig 1B to 1D).

**Figure 1.**
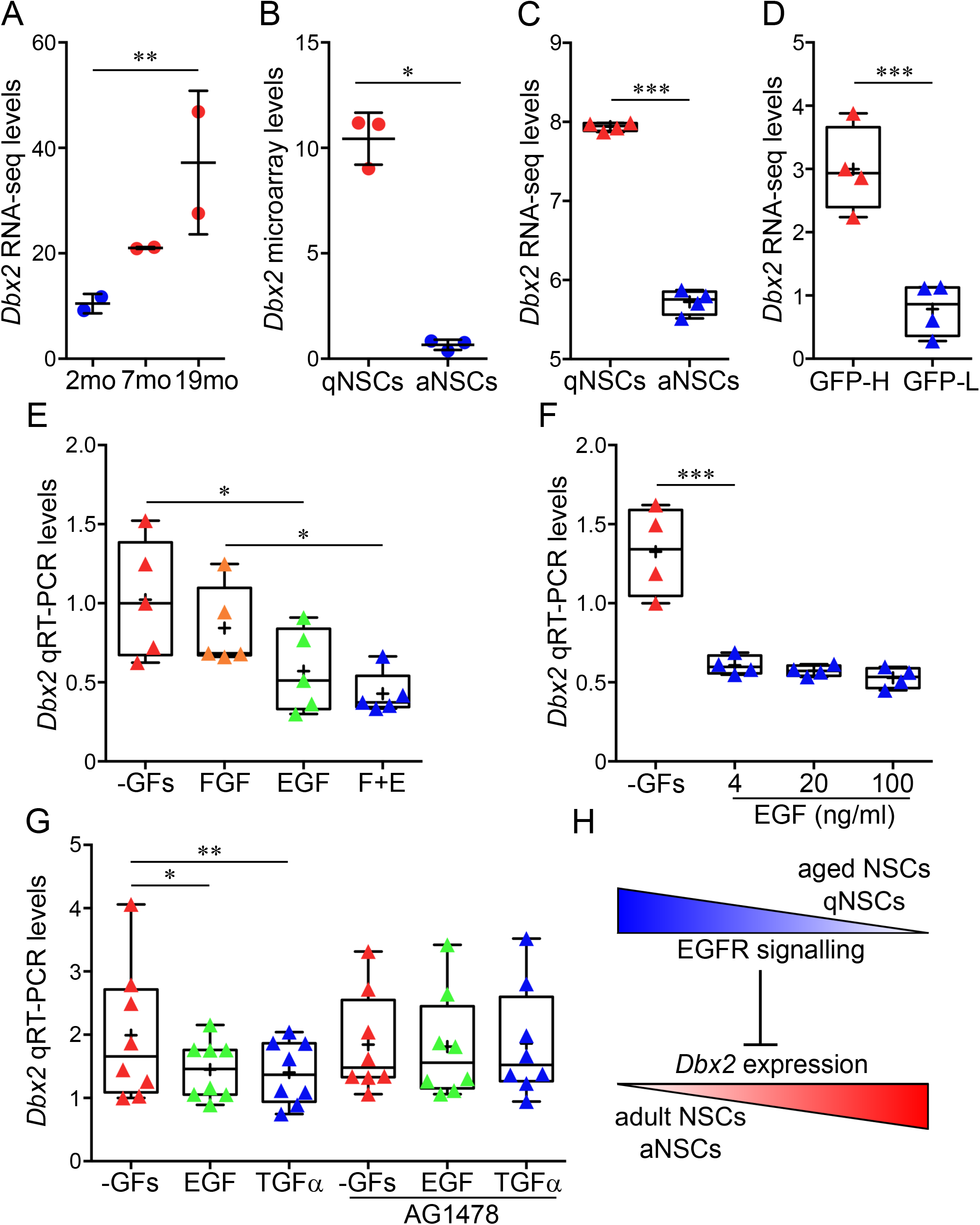
Activation and EGFR signaling downregulate *Dbx2* expression in adult NSPCs. **(A)** *Dbx2* mRNA levels in freshly sorted NSPCs from the SVZ of 2 mo, 7 mo and 19 mo mice as reported in a published (Kalamakis *et al*, 2019) RNA-seq dataset. Black lines show mean transcript levels ± standard deviation; blue and red dots show individual experimental replicates for young adult and aged mice, respectively (n=2); **, adjusted p value < 0.01, 2 mo vs 19 mo NSPCs. **(B)** *Dbx2* mRNA levels in freshly sorted qNSCs (red dots) and aNSCs (blue dots) from the SVZ of adult mice as reported in a published (Codega *et al*, 2014) microarray dataset (n=3); *, adjusted p value < 0.05. **(C)** Box-and-whisker plots of *Dbx2* mRNA levels in freshly sorted qNSCs and aNSCs from the SVZ of adult mice as reported in a published (Belenguer *et al*, 2021) RNA-seq dataset. The lower and higher whiskers indicate the minimum and maximum values, respectively. The bottom and top of the box represent the first and third quartiles, respectively, and the band inside the box indicates the second quartile (the median). The + symbol indicates mean transcript levels. Red and blue triangles show individual experimental replicates for qNSCs and aNSCs, respectively (n=4); ***, p < 0.001, Student’s t-test. **(D)** Box-and-whisker plots of *Dbx2* mRNA levels in freshly sorted GFP high (GFP-H, red triangles) and GFP low (GFP-L, blue triangles) NSCs (largely corresponding to qNSCs and aNSCs, respectively) from the SVZ of adult mice as reported in a published (Xie *et al*, 2020) RNA-seq dataset (n=4); ***, adjusted p value < 0.001. **(E)** Box-and-whisker plots of *Dbx2* mRNA levels in adherent cultures of mouse young adult SVZ NSPCs that were treated for 6h without exogenous growth factors (-GFs, red triangles), with 10 ng/ml FGF2 (orange triangles), with 20 ng/ml EGF (green triangles), or with EGF + FGF2 (E+F, blue triangles), followed by qRT-PCR analysis (n=5); *, p < 0.05, one-way ANOVA. **(F)** Box-and-whisker plots of *Dbx2* mRNA levels in NSPC cultures that were treated for 8h without exogenous growth factors (-GFs, red triangles), or were maintained for 6h without growth factors and then treated for 2h with 4, 20 or 100 ng/ml EGF (blue triangles), followed by qRT-PCR analysis (n=4); ***, p < 0.001, one-way ANOVA. **(G)** Box-and-whisker plots of *Dbx2* mRNA levels, as detected by qRT-PCR, in NSPC cultures that were treated for 6h without exogenous growth factors (-GFs, red triangles), with 20 ng/ml EGF (green triangles), or with 100 ng/ml TGFα (blue triangles), in the absence or in the presence of 2 μM AG1478 (n=8); **, p < 0.01, one-way ANOVA. **(H)** Proposed model of *Dbx2* regulation in adult SVZ NSPCs. EGFR signaling inhibits Dbx2 expression. EGFR signaling levels are reduced in qNSCs and in aged NSCs in comparison with aNSCs and adult NSCs (blue gradient), leading to increased Dbx2 expression during NSC quiescence or aging (red gradient).

We then investigated whether EGFR signaling may be involved in *Dbx2* expression changes upon NSC activation or NSPC aging. To this goal, we analysed *Dbx2* mRNA levels in cultures of young adult mouse SVZ NSPCs that were maintained for 6 hours (6h) in standard proliferation-supporting media, supplemented with EGF and FGF2, in media devoid of both growth factors, or in media containing only EGF or FGF2. As shown in Fig 1E, *Dbx2* was upregulated in NSPC cultures devoid of exogenous EGF. Furthermore, *Dbx2* was downregulated when NSPCs cultured without growth factors for 6h were then treated with EGF for 2h (Fig 1F). TGFα is the predominant EGFR ligand in the adult brain and its levels decrease in the aged SVZ (Enwere *et al*, 2004). We observed a comparable decrease in *Dbx2* mRNA levels in NSPC cultures treated for 6h with EGF or TGFα, in comparison with NSPCs cultured without exogenous growth factors; this decrease was prevented by including AG1478, an EGFR inhibitor, in the culture media (Fig 1G). These results suggest that EGFR signaling represses *Dbx2* expression in SVZ NSPCs; consequently, a reduction of EGFR activation during quiescence or aging may lead to *Dbx2* upregulation in NSPCs (Fig 1H).

### *Dbx2* overexpression in young adult mouse SVZ NSPCs inhibits cell proliferation by hindering the G2/M transition

*Dbx2* overexpression reduces the growth of NSPC cultures derived from the SVZ of young adult mice, without major effects on NSPC viability, suggesting that *Dbx2* inhibits NSPC proliferation (Lupo *et al*, 2018). We speculated that *Dbx2* may promote cell cycle exit and entry in G0, or decrease the division rate of NSPCs remaining in the cell cycle. To test this hypothesis, we took advantage of previously described transgenic young adult NSPC lines with constitutive expression of either a *Dbx2* transgene (*Dbx2*-NSPCs) or a control *GFP* transgene (*GFP*-NSPCs), and cultured them for 4-6 days (4-6d) in non-adherent conditions to generate neurospheres (Lupo *et al*, 2018). We then used these cultures to quantify the percentage of *Dbx2*-NSPCs and *GFP*-NSPCs that were immunostained for Ki67, a nuclear marker expressed throughout the cell cycle, but not in cells that have exited the cell cycle. Moreover, we estimated the fraction of *Dbx2*-NSPCs and *GFP*-NSPCs in each phase of the cell cycle (G0/G1, S, G2/M) by flow cytometry quantification of DNA content in cells stained with propidium iodide (PI). To make sure that the results were reproducible, we performed these analyses with two different pairs of *Dbx2*-NSPC and *GFP*-NSPC lines, obtained from independent derivations of young adult mouse SVZ NSPCs (Lupo *et al*, 2018). Cell cycle exit of NSPCs in response to *Dbx2* overexpression would be expected to cause an increase in the fraction of cells in G0/G1 and a decrease of the fraction of cells positive for Ki67; however, these effects were not detectable in the *Dbx2*-NSPC lines as compared to *GFP*-NSPC cultures (Fig 2A and 2E; Fig EV1A and EV1E; Appendix Fig S1A and S1D; representative images of Ki67-stained cultures are shown in Fig 2I and 2J, and in Fig EV1I and EV1J). Surprisingly, we detected a reproducible increase of the G2/M cell fraction upon *Dbx2* overexpression (Fig 2C; Fig EV1C). To distinguish between cells in G2 and in M phases, we quantified the fraction of cells immunostained for phosphorylated histone H3 (pH3), a marker of cells undergoing mitosis. Mitotic histone H3 phosphorylation starts in pericentric heterochromatin in late G2, then spreads throughout the condensing chromatin during prophase, reaching a peak at metaphase; dephosphorylation of histone H3 begins in anaphase and ends in telophase (Hendzel *et al*, 1997). In agreement with the dynamics of mitotic pH3 accumulation, we observed different patterns of pH3 immunostaining during NSPC progression through mitosis. Some of the pH3-positive (pH3+) NSPCs, likely corresponding to late G2 cells, showed a weaker staining in the form of isolated nuclear spots; during prophase, these pH3+ foci increased in size and intensity, until the whole nucleus appeared to be stained (Appendix Fig S2A to S2D). A peak in staining intensity was observed in NSPC nuclei between metaphase and early anaphase, followed by a progressive decrease between anaphase and telophase (Appendix Fig S2I to S2L). We quantified the total percentage of pH3+ NSPCs, which included all the cells engaging in the G2/M transition, as well as the percentage of NSPCs showing pH3 staining patterns typical of cells that have progressed beyond early prophase (Appendix Fig S2D and S2I to S2L), which we named pH3+ late cells. When compared with *GFP*-NSPCs, a decrease in the pH3+ total cell fraction was detected in one of the *Dbx2*-NSPC lines, but not the other; however, both lines showed a reduction in the pH3+ late cell fraction (Fig 2F and 2G; Fig EV1F and EV1G; Appendix Fig S2B and S2E; representative images of pH3-stained cultures are shown in Fig 2K and 2L, and in Fig EV1K and EV1L). The fraction of pH3+ cells within the top 40% of the fluorescence intensity range, which likely include pH3+ late cells, was also reduced in both *Dbx2*-NSPC lines (Appendix Fig S2C and S2F). These results indicate that the accumulation in G2/M observed by flow cytometry is due to an altered mitotic progression in *Dbx2*-NSPC cultures. Therefore, they suggest that *Dbx2* negatively regulates the G2/M transition in NSPCs, hindering the progression through mitosis when its expression levels increase. The fact that one of the *Dbx2*-NSPC lines showed a weaker effect on the G2/M transition than the other is consistent with the comparatively weaker growth phenotype of the neurosphere cultures generated from this cell line (Lupo *et al*, 2018). Nonetheless, the number of cells undergoing cell division is decreased in both lines, which provides an explanation for the reduced size of the neurospheres formed by *Dbx2*-NSPCs.

**Figure 2.**
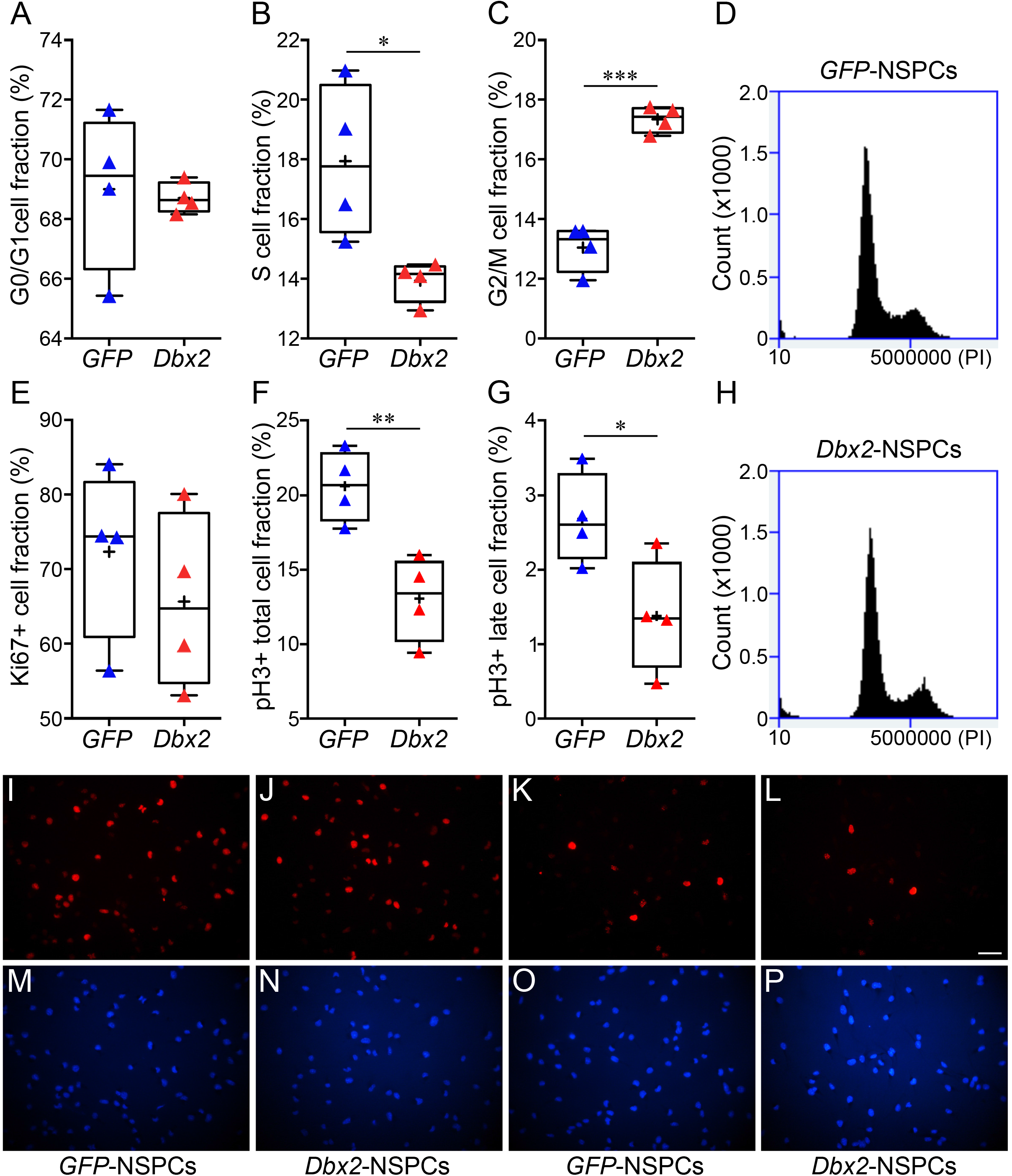
Constitutive *Dbx2* overexpression inhibits the G2/M transition in young adult NSPCs. **(A** to **C)** Box-and-whisker plots of the fraction of *GFP*-NSPCs (blue triangles) and *Dbx2*-NSPCs (red triangles) in the G0/G1 **(A)**, S **(B)** and G2/M **(C)** phases of the cell cycle (n=4); *, p < 0.05, ***, p < 0.001, Student’s t-test. Flow cytometry histograms of PI stained *GFP*-NSPC and *Dbx2*-NSPC cultures from a representative experiment are shown in **(D)** and **(H)**, respectively. **(E** to **G)** Box-and-whisker plots of the fraction of Ki67+ **(E)**, pH3+ total **(F)** and pH3+ late **(G)** cells in *GFP*-NSPC (blue triangles) and *Dbx2*-NSPC (red triangles) cultures (n=4); *, p < 0.05, **, p < 0.01, Student’s t-test. **(I** to **P)** Representative images of *GFP*-NSPC **(I**, **M**, **K**, **O)** and *Dbx2*-NSPC **(J**, **N**, **L**, **P)** cultures stained with anti-Ki67 **(I**, **J)** or anti-pH3 **(K**, **L)** antibodies. Hoechst nuclear staining is shown in **(M** to **P)**. Scale bar, 40 μm.

### *Dbx2* regulates NSPC transcriptional programmes associated with cell cycle, inflammatory response and lineage progression

To gain insight into the molecular mechanisms mediating the effects of *Dbx2* overexpression in SVZ NSPCs, we performed a transcriptomic analysis with the transgenic NSPC lines used for cell cycle assays. To this aim, we performed RNA-sequencing (RNA-seq) and differential gene expression analysis with RNA samples obtained from 4-6d non-adherent cultures of both lines of *Dbx2*-NSPCs and *GFP* NSPCs (4 independent experiments for each pair). Principal component analysis (PCA) confirmed that the cell culture samples used for transcriptomic analysis were mainly clustered according to transgene expression (*Dbx2* or *GFP*) (PC1, 63% of variance), although samples belonging to the same experimental condition, but to independent cell lines, could also be distinguished (PC2, 21% of variance) (Fig 3C). By setting a threshold of false discovery rate (FDR) < 0.05, we identified 2854 differentially expressed genes (DEGs) between *Dbx2*-NSPCs and *GFP*-NSCPs (1368 upregulated and 1486 downregulated genes in *Dbx2*-NSPCs) (Fig 3A and Table S1).

**Figure 3.**
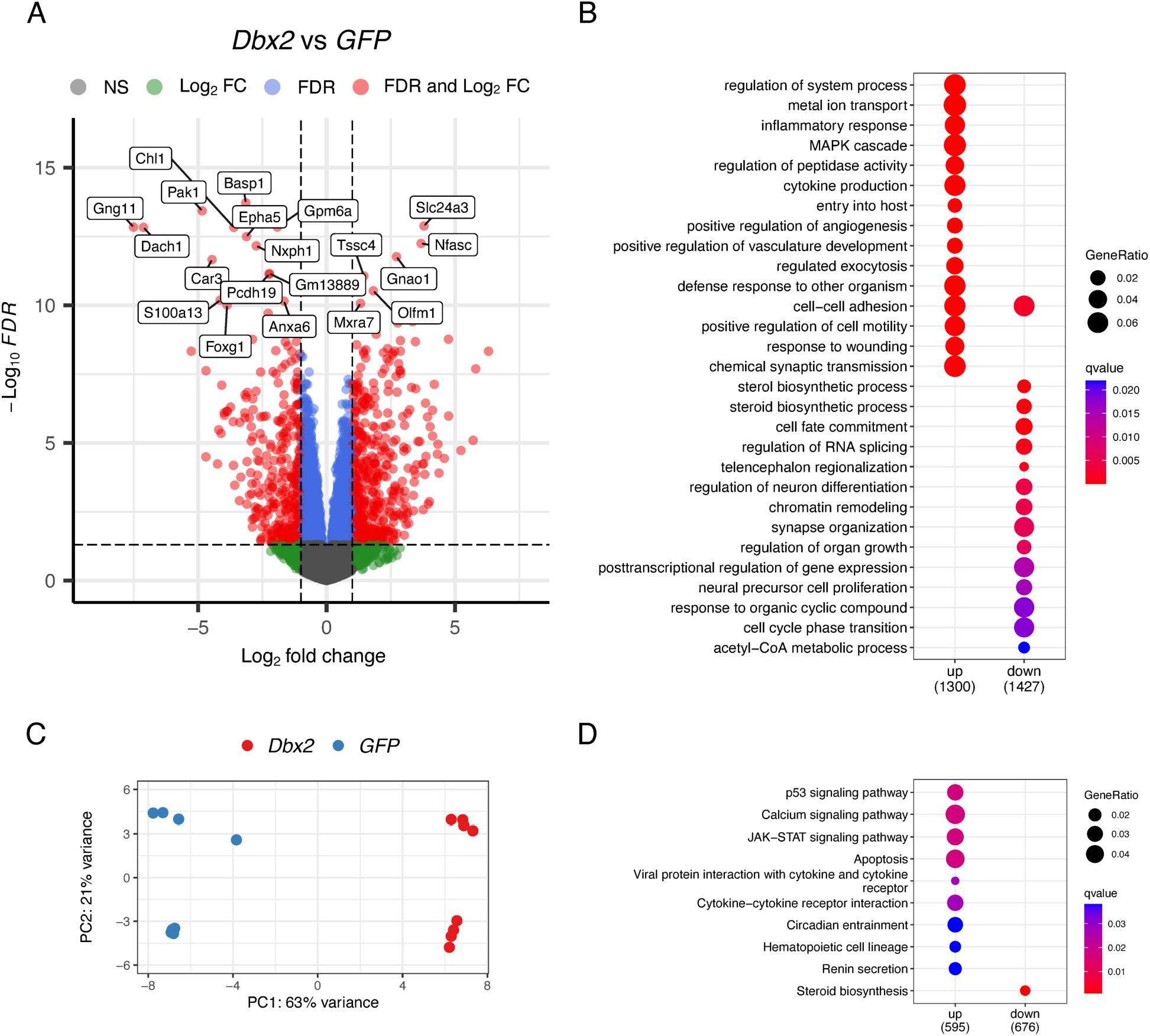
Transcriptomic profiling of *Dbx2*-overexpressing NSPCs. **(A)** Volcano plot showing the distribution of −log_10_(FDR) values (Y axis) relative to log_2_(FC) values (X axis) resulting from the comparison of gene expression levels in *Dbx2*-NSPCs vs *GFP*-NSPCs. Red dots indicate DEGs associated with FDR < 0.05 and FC < −2 or FC >2; blue dots indicate genes with FDR < 0.05 and −2 < FC < 2; green dots indicate genes with FC < −2 or FC >2, and FDR > 0.05. Representative genes are highlighted in the plot. **(B)** Dot plot showing the GO terms enriched in the upregulated (left) and the downregulated (right) DEGs. The size of the dots is based on the count of the DEGs correlated with each term; the colour of the dots shows the qvalue associated with gene enrichment for each term. **(C)** PCA analysis plots showing that control experimental replicates (*GFP*-NSPCs, blue dots) and *Dbx2*-overexpressing experimental replicates (*Dbx2*-NSPCs, red dots) can be distinguished from each other along the PC1 axis. Control or *Dbx2*-overexpressing experimental replicates from independent cell lines carrying the same transgene can be distinguished along the PC2 axis. **(D)** Dot plot showing the KEGG pathways that are enriched for the upregulated (left) and the downregulated (right) DEGs.

We then performed a gene ontology (GO) enrichment analysis of the DEGs using the GO Biological Process database. As shown in Fig 3B, the genes downregulated upon *Dbx2* overexpression were enriched for GO terms related to NSPC proliferation (e.g. “neural precursor cell proliferation”, “cell cycle phase transition”), lineage progression (e.g. “cell fate commitment”, “regulation of neuron differentiation”), and RNA processing (e.g. “RNA splicing”); the genes upregulated in *Dbx2*-overexpressing NSPCs were enriched for GO terms related to inflammatory response (e.g. “inflammatory response”, “cytokine production”). DEG analysis using REACTOME and KEGG databases also indicated that pathways related to cell cycle and RNA processing were enriched in the downregulated genes, and pathways related to inflammatory response and cell cycle inhibition (“p53 signaling pathway”) were enriched in the upregulated genes (Fig 3D and Appendix Fig S3).

To characterise these transcriptional effects in more detail, we analysed the transcriptome of *Dbx2*-overexpressing NSPCs by gene set enrichment analysis (GSEA), a computational method that investigates the biological effects associated with a ranked gene list by determining whether the members of a gene set linked to a specific biological process tend to occur at the top or the bottom of the ranked gene list (Subramanian *et al*, 2005). Since the size of the analysed gene list influences GSEA accuracy, we performed this analysis with a wider *Dbx2*-associated signature of 12533 genes, which included all the genes with detectable mRNA expression in *Dbx2*-NSPCs and GFP-NSPCs, irrespectively of FDR values. This gene list was ranked according to fold change (FC), followed by GSEA with “Hallmark” gene sets in the Molecular Signatures Database (MSigDB). Notably, several cell cycle-related gene sets (e.g. “E2F targets”, “G2M checkpoint”, “Myc targets”) correlated with the *Dbx2*-associated signature (Fig 4A), with negative NES values resulting from the enrichment of these gene sets among the genes downregulated in *Dbx2*-overexpressing NSPCs (Fig 4B, 4C and 4D; Table S2). Moreover, several gene sets linked to inflammatory response (e.g. “Interferon gamma response”, “IL6 JAK STAT3 signaling”) or to cell cycle inhibition (“p53 pathway”) correlated with the *Dbx2*-associated signature (Fig 4A), with positive NES values resulting from the enrichment of these gene sets among the genes upregulated in *Dbx2*-overexpressing NSPCs (Fig 4E, 4F and 4G, Table S2). Transcription factor motif enrichment analysis confirmed an enrichment of E2F and Myc motifs in the DEGs downregulated in *Dbx2*-overexpressing NSPCs, and an enrichment of p53 motifs in the upregulated DEGs (Fig EV24).

**Figure 4.**
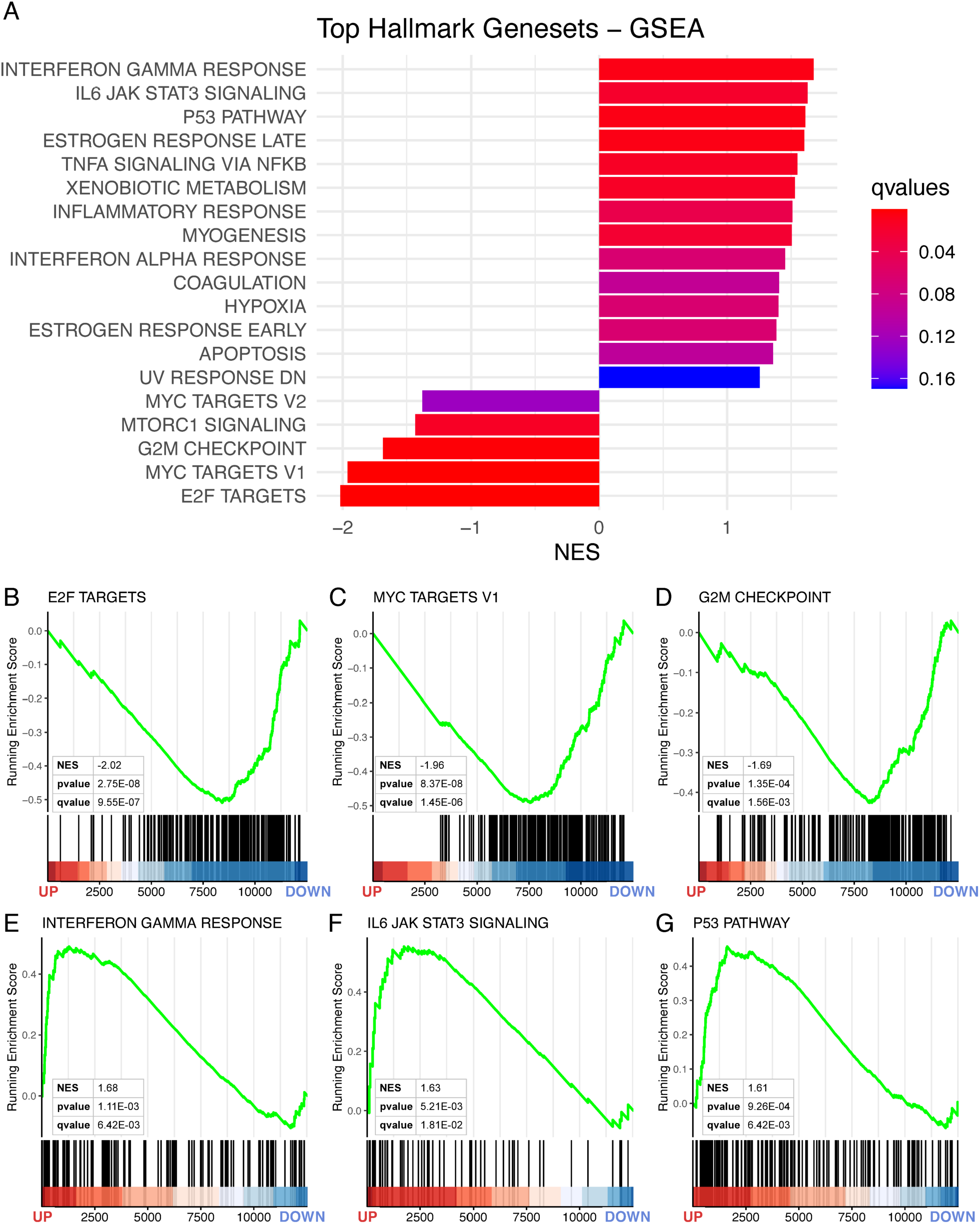
*Dbx2* overexpression affects the expression of gene sets related to cell cycle and inflammatory response. **(A)** Bar plot showing the top “Hallmark” gene sets correlated with the *Dbx2*-associated transcriptional signature (12533 genes), according to GSEA. The size and the colour of the bars indicate the NES value and the qvalue associated with each gene set, respectively. (**B** to **G**) Enrichment plots showing the top 3 NES < 0 gene sets (**B** to **D**), or the top 3 NES > 0 gene sets (**E** to **G**), in the *Dbx2*-associated signature. Vertical black lines indicate individual members of each gene set; the heat maps at the bottom of the plots show the position of each gene within the ranked *Dbx2*-associated signature.

Given *Dbx2* effects on NSPC proliferation, we performed a more detailed analysis of the transcriptional changes related to cell cycle progression in *Dbx2*-overexpressing NSPCs. To this aim, we carried out GSEA using the *Dbx2*-associated signature and several gene sets related to different cell cycle phases as available in the “C2” curated gene sets collection in MSigDB. This analysis revealed a general negative correlation of the *Dbx2*-associated signature with cell cycle-related gene sets (Fig EV3). A characterization of the DEGs between *Dbx2*-NSPCs and *GFP*-NSCPs (FDR < 0.05) with KEGG cell cycle and p53 signaling pathway gene sets showed that many genes crucially implicated in the G1/S transition (e.g. *Cdk2, Cdk4, Cdk6*) or G2/M transition (e.g. *CycA, Cdk1*) were downregulated in *Dbx2*-overexpressing NSPCs, and key cell cycle inhibitors acting in the p53 pathway (e.g. *p21, Gadd45*) were upregulated (Fig EV4). These results suggest that elevated *Dbx2* expression broadly inhibits the transcriptional pathways involved in cell cycle progression.

The results of GSEA with cell cycle-related gene sets prompted us to use the same approach to investigate the effects of *Dbx2* overexpression on the transcriptional programmes associated with NSPC lineage progression. To this aim, we took advantage of the recently described single-cell transcriptomic signatures of qNSCs, aNSCs, TAPs and NBs of the adult mouse SVZ (Zywitza *et al*, 2018)(Kalamakis *et al*, 2019)(Xie *et al*, 2020); we then used the gene sets related to each of these cell populations to perform GSEA with the *Dbx2*-associated signature. Remarkably, the gene sets related to aNSCs, TAPs and NBs negatively correlated with the *Dbx2*-associated signature (Fig 5, Table S3). Although the signature related to the whole qNSC population was not correlated with the *Dbx2*-associated signature (Fig 6A), a positive correlation (NES > 0) was observed when the analysis was carried out using the gene sets related to specific qNSC subpopulations (Fig 5C and 5E). Since aNSCs, TAPs and NBs encompass the proliferating NSPC pool of the SVZ neurogenic lineage, we wondered whether the negative correlation that we observed between the gene sets related to these cell populations and the *Dbx2*-associated signature may be driven by cell cycle-related genes. To address this question, we quantified the percentage of REACTOME and KEGG cell cycle genes among the aNSC-related and TAP-related gene sets. This quantification suggested that cell cycle genes alone did not explain the correlation between the aNSC/TAP-related and the *Dbx2*-associated signatures; for example, cell cycle genes accounted for only 26-28% of the genes in the signatures related to proliferative TAPs (mTAPs and L1) (Fig 5B, 5D and 5F). We then filtered out cell cycle genes from aNSC/TAP signatures, selected the *Dbx2*-modulated DEGs (FDR < 0.05) among the remaining genes, and performed GO enrichment analysis on the resulting gene lists. Notably, GO terms related to RNA processing (e.g. “RNA splicing”) prominently featured in all the analysed gene lists (Fig EV5). These data suggest that *Dbx2* may inhibit NSPC proliferation by regulating cell cycle-related genes, and NSPC lineage progression by modulating genes involved in RNA processing.

**Figure 5.**
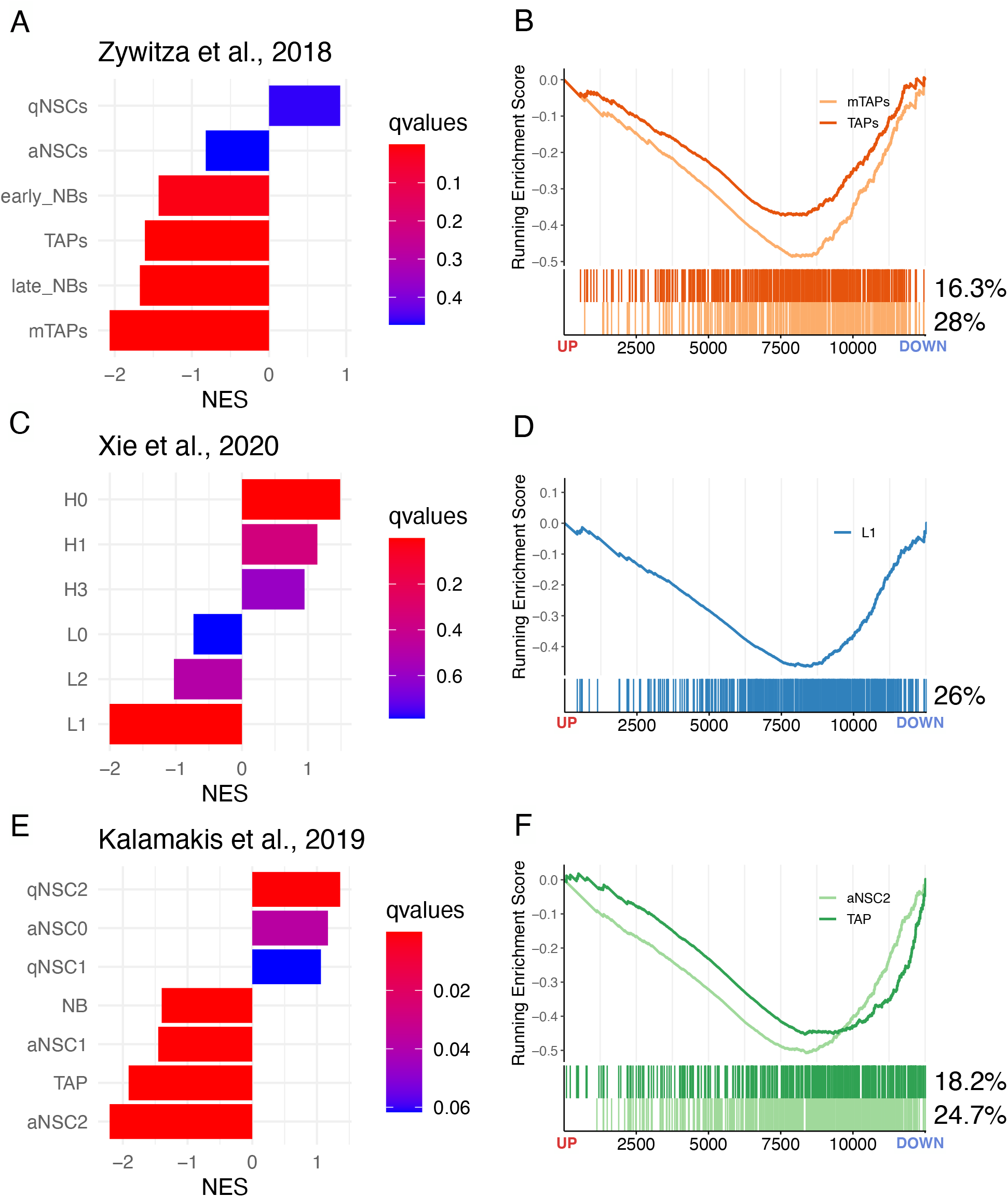
*Dbx2* affects the expression of gene sets related to neurogenic lineage progression. **(A**, **C** and **E)** Bar plots showing the correlation between the *Dbx2*-associated signature and the qNSC-specific, aNSC-specific, TAP-specific and NB-specific gene sets, as defined in recent studies [(Zywitza *et al*, 2018) **(A)**, (Xie *et al*, 2020) **(C)**, (Kalamakis *et al*, 2019) **(E)**], according to GSEA. In **(C)**, H0-H3 correspond to qNSC subpopulations, L0 to aNSCs, L1 to TAPs and L2 to NBs, as previously described (Xie *et al*, 2020). (**B**, **D** and **F**) Enrichment plots showing selected cell type-specific gene sets in the *Dbx2*-associated signature. The FC direction in *Dbx2*-overexpressing NSPCs is indicated at the bottom of the plots (down, FC <0; red, FC >0). The percentages indicate the fraction of cell cycle genes in each gene set.

In this study, we show that elevated *Dbx2* expression in NSPCs impair the G2/M transition, causing accumulation in G2 by hindering the progression through mitosis. Although we could not detect a clear effect of *Dbx2* overexpression on the G1/S transition, transcriptomic analyses revealed that *Dbx2* broadly regulates the cell cycle molecular networks, repressing genes implicated in cell cycle progression and activating genes involved in cell cycle arrest. Of note, signatures related to E2F, Myc and p53 were among the top hits, which is consistent with the recently described association between these factors and the transcriptomic profiles of aged NSPCs (Xie *et al*, 2020).

Recent studies suggest that the increase in the time spent by NSCs in a quiescent state, which is generally thought to be in G0, is a main driver of the neurogenic decline of aged mice. Although *Drosophila* qNSCs can also be arrested in G2 (Otsuki & Brand, 2018), evidence that this may happen in mice is lacking. Transcriptomic comparison of qNSCs and aNSCs of the mouse SVZ, however, showed that both G1/S and G2/M signatures are downregulated in qNSCs (Belenguer *et al*, 2021); thus, the inhibitory effects of *Dbx2* on the transcriptional networks associated with both the G1/S and G2/M transitions is compatible with a role in the increased quiescence of aged NSCs. Notably, the gene set downregulated upon *Dbx2* overexpression correlates with the transcriptomic signatures of the proliferative populations of the SVZ neurogenic lineage (aNSCs, TAPs and early NBs), supporting *Dbx2* implication in NSC quiescence from a different angle. The limited percentage of cell cycle genes contributing to this correlation suggests that it is not mainly driven by cell cycle inhibition; furthermore, GO analysis indicates an additional negative effect of *Dbx2* on genes related to RNA processing. We speculate that increased *Dbx2* expression during SVZ aging may facilitate NSC sliding into quiescence by inhibiting cell cycle genes along with RNA processing genes.

Several categories associated with inflammatory response were top hits among the genes upregulated by *Dbx2* overexpression. This is noteworthy, since inflammatory transcriptional signatures are strongly associated with aged neurogenic niches and brain aging (Lupo *et al*, 2019a)(Lupo *et al*, 2019b)(Navarro Negredo *et al*, 2020). We did not detect an increase in *Dbx2* expression levels in young adult NSPC cultures treated with IFNαγ, IL2/6 or TNFα (our unpublished observations); however, gene sets related to immune response (especially IFN signaling) were upregulated in SVZ NSPC cultures treated with BMP4 and FGF2 to induce quiescence (Marqués-Torrejón *et al*, 2021). Thus, these signatures may be activated in NSPCs independently of canonical cytokine-driven responses to regulate their proliferative state.

In conclusion, our study uncovers several links between increased *Dbx2* function and key features of NSPC aging. These include: i) the downregulation of gene networks implicated in cell cycle progression, the upregulation of those involved in cell cycle arrest and the inhibition of NSPC proliferation; ii) the upregulation of inflammatory response pathways; iii) the downregulation of the transcriptomic signatures of the proliferative neurogenic cell populations (aNSCs, TAPs and early NBs), which are depleted in the aged SVZ; iv) the negative regulation of *Dbx2* expression by EGFR signaling, which is reduced in the aged SVZ. This work has obvious limitations, since our functional experiments have been performed using *in vitro* NPSC cultures and gain-of-function approaches. Although some of the genes modulated upon *Dbx2* overexpression showed opposite changes in aged NSPC cultures expressing *Dbx2*-targeting shRNAs (our unpublished observations), this approach yielded a modest *Dbx2* knockdown in our hands (Lupo *et al*, 2018). Of note, *Dbx2* is a NSC-specific component of the recently reported molecular aging clocks in the mouse SVZ, and its age-associated modulation can be reversed by exercise (Buckley *et al*, 2022). Gene function analyses *in vivo* by conditional genetic manipulation in adult NSPCs are expensive and time consuming; our *in vitro* analyses do not conclusively demonstrate a functional role of *Dbx2* in neurogenic aging but provide a strong case for future studies investigating this gene in the context of *in vivo* neurogenic niches.

## MATERIALS AND METHODS

### NSPC culture

This work was carried out by *in vitro* culture of available liquid nitrogen stocks of mouse NSPCs that were previously derived from the SVZ of 3 months old (3 mo) young adult mice. The original derivation of mouse SVZ NSPCs was performed in accordance with EU and Italian regulations and with ethical approval by the Ethical Committee for Animal Research of the Italian Ministry of Health, as described (Lupo *et al*, 2018). No additional animals were employed for the experiments reported in the present study. NSPC culture in adherent and non-adherent proliferating conditions using media supplemented with human recombinant EGF and FGF2 (R&D Systems) was performed according to published protocols (Soldati *et al*, 2012)(Carucci *et al*, 2017)(Lupo *et al*, 2018). Transgenic NSPC lines with constitutive expression of a *Dbx2* transgene or a control *GFP* transgene were previously described (Lupo *et al*, 2018), and were used again for this study.

### Real-time qRT-PCR assays

For the analysis of *Dbx2* mRNA levels by real-time qRT-PCR, young adult SVZ NSPCs were seeded in coated T25 flasks or 6-well plates at a density of approximately 40000 cells/cm^2^ in media for adherent proliferative culture conditions. On the next day, cultures were rinsed once with basal media devoid of growth factors, then incubated in fresh media with different combinations of growth factors as described in the Results section. Treatments with human recombinant Transforming Growth Factor α (TGFα, Cell Guidance Systems) were performed by diluting a 50 μg/ml stock in PBS containing 0.1% Bovine Serum Albumine with culture media. Treatments with the EGFR inhibitor AG1478 (Calbiochem) were performed by diluting a 10 mM stock in DMSO with culture media; control cultures were treated with equal volumes of DMSO. At the end of the experimental treatment, cells cultured in T25 flasks were enzimatically detached, pelleted in 2 ml tubes and lysed for RNA purification, whereas cells in 6-well plates were directly lysed in the culture plates. Total RNA purification and quantification, reverse transcription and real-time qPCR were performed using Qiagen kits, a NanoDrop 2000 (Thermo Scientific) and a Rotor-Gene Q (Qiagen), as previously described (Carucci *et al*, 2017). Primer sequences for *Dbx2* and *Rpl19*, used as reference gene, were previously reported (Lupo *et al*, 2018).

### Cell proliferation analyses by flow cytometry and immunocytochemistry

For cell cycle analysis, neurospheres generated by non-adherent culture of control or *Dbx2*-overexpressing NSPCs were pelleted by centrifugation and dissociated with Accutase (Corning). The resulting cell suspensions were rinsed with PBS, fixed in cold PBS-methanol 1:1 and stored in this solution at 4°C. Staining with PI and quantification of the fraction of cells in G0/G1, S and G2/M by flow cytometry were performed as previously described (Licursi *et al*, 2020).

To quantify the fraction of cells expressing Ki67 and pH3, cells suspensions obtained by centrifugation and dissociation of control and *Dbx2*-overexpressing neurospheres were plated on poly-ornithine/laminin coated glass coverslips in 24-well plates and cultured for 2-4 hours in media for adherent proliferative conditions to allow attachment to the glass. After attachment, cells were fixed for 20 min with methanol-free formaldehyde (Pierce) diluted to 4% with PBS, then rinsed a few times with PBS and stored in PBS at 4°C. Immunofluorescence analysis with an anti-Ki67 mouse monoclonal antibody (Leica Biosystems, 1:100) or with an anti-pH3 (Ser 10) rabbit polyclonal antibody (ThermoFisher, 1:100) was performed as previously described (Licursi et al., 2020; Lupo et al., 2018). In each experiment, 12 random fields per coverslip were usually selected based on Hoechst staining and photographed using a Nikon Eclipse TE300 microscope and a Nikon DS-U3 digital camera, followed by the quantification of the ratio between Ki67+ or pH3+ cells and the total cell number in 1-3 coverslips for each experimental condition. Several hundred to a few thousand cells were counted for each experimental condition in each experiment. Automated immunofluorescence analysis was performed with ImageJ software (Schneider *et al*, 2012). Colocalization images were generated using the ImageJ plug-in “Colocalization Image Creator” (Lunde & Glover, 2020). For Hoechst signal, the “Binary Element” option was used and the automatic thresholding was set on “YEN”; for Ki67 and pH3 signals, the “Grayscale Element” option was chosen. The colocalization images generated were then used for automated counting of Ki67+ or pH3+ cells using the ImageJ plug-in “Colocalization Object Counter” (Lunde & Glover, 2020). Only Ki67 and pH3 signals effectively colocalizing with Hoechst signal were counted. Automated analysis of signal intensity was performed using the ImageJ tool “Analyse Particles”. A nuclear mask was generated using the Hoechst signal to specifically select nuclear regions. This mask was then merged with the Ki67 or pH3 signals and the fluorescence intensity per pixel in each nucleus was measured with the ImageJ tool “Measure”.

### Statistical analysis of gene expression and cell proliferation assays

The experimental data obtained from qRT-PCR, flow cytometry and immunofluorescence assays were analysed and graphically represented using GraphPad Prism 9 software. The type of statistical test performed, the resulting p values and the number of independent experimental replicates performed with NSPCs from different culture passages (n) are indicated in the figure legends.

### RNA-seq and differential gene expression analysis

For RNA-seq, total RNA was purified from frozen cell pellets of control and *Dbx2*-overexpressing non-adherent cultures using Qiagen RNeasy kits. RNA-seq libraries from total RNA (100 ng) from each sample were prepared using QuantSeq 3’ mRNA-Seq Library prep kit (Lexogen, Vienna, Austria), according to manufacturer’s instructions, at Next Generation Diagnostics (Pozzuoli, Italy). The amplified fragmented cDNAs of 300 bp in size were sequenced in single-end mode using the NextSeq500 (Illumina) with a read length of 101 bp. Sequence read quality was evaluated using *FastQC* version 0.11.8 (Babraham Institute, Cambridge, UK), followed by trimming using *bbduk* software to remove adapter sequences, poly-A tails and low-quality end bases (Q < 20). Reads were then mapped to the mouse Ensembl GRCm38 (mm10) build reference genome with *STAR* version 2.5.0a (Dobin *et al*, 2013); gene annotations corresponding to Ensembl annotation release 96 were used to build a transcriptome index, which was provided to *STAR* during the alignment.

To identify DEGs, data were filtered to remove from the analysis the genes having < 1 counts per million in less than 7 out of 16 total samples for each comparison. Data normalization and differential gene expression analysis were performed using Bioconductor, R package *edgeR* version 3.36 (Robinson *et al*, 2010), assigning the cell line batch as a covariate to fit the generalized log-linear model with the glmQLFit function of *edgeR*. DEGs were assessed by comparing *Dbx2*-NSPC and *GFP*-NSPC samples using a moderated t-test with a FDR threshold < 0.05. Volcano plot was created using Bioconductor (Gentleman *et al*, 2004)(Huber *et al*, 2015), R package *EnhancedVolcano* version 1.12.0. PCA was performed using R base prcomp function, plotting the first two PCs using R package *ggplot2* version 3.3.5.

### Gene ontology, GSEAand transcription factor enrichment analyses

DEGs in *Dbx2*-NSPCs vs *GFP*-NSPCs were clustered by functional annotation in GO and pathway enrichment analysis using Bioconductor R package *clusterProfiler* version 4.2.0 (Wu et al., 2021) with annotation of Gene Ontology Database (Ashburner *et al*, 2000) and with annotation of REACTOME (https://doi.org/10.1093/nar/gkt1102) and Kyoto Encyclopedia of Genes and Genomes (KEGG) (Ogata *et al*, 1999) for pathways. R package *pathview* version 1.34.0 was used to integrate the RNA-seq data with KEGG pathway plots.

BROAD Institute GSEA (Subramanian *et al*, 2005) was used to assess the enrichment of the *Dbx2*-associated signature list (12533 expressed genes in Dbx2-NSPCs and GFP-NSPCs without FDR threshold), ranked according to FC, versus the curated “Hallmark” and “C2” gene set collections from the BROAD Molecular Signatures Database version 7.4.1 (https://www.gsea-msigdb.org/gsea/msigdb/), or versus recently published mouse SVZ single-cell RNA-seq datasets (Zywitza *et al*, 2018)(Kalamakis *et al*, 2019)(Xie *et al*, 2020). The mouse version of the gene set collections “Hallmark” and “C2” (REACTOME subcategory, selecting signatures related to cell cycle) were obtained from R package *msigdbr* version 7.4.1. GSEA enrichment score (ES) was calculated by walking down the ranked list of genes, increasing a running-sum statistic when a gene was in the gene set and decreasing it when it was not. A normalized ES (NES) was also calculated with GSEA, by considering differences in pathway size (i.e., gene set size) and allowing for comparisons between pathways within the analysis.

Transcription factor enrichment analysis was performed with Enrichr method (https://doi.org/10.1093/nar/gkw377), using as references TRANSFAC and JASPAR (https://doi.org/10.1093/nar/gkab1113) PWMs library of transcription factors, containing manually curated transcription factors binding profiles as position weight matrices (PWMs), TRRUST (https://doi.org/10.1093/nar/gkx1013) library, a database of reference transcription factor–target regulatory interactions, and ENCODE and CHEA library, containing experimental transcription factor binding data from ENCODE Project Consortium (https://doi.org/10.1038/s41586-020-2493-4) and CHEA database (https://doi.org/10.1093/bioinformatics/btq466).

## ACKNOWLEDGMENTS

We thank Daniela Trisciuoglio and Paola Del Porto for help with the flow cytometry data shown in Fig 2 and Fig EV1.

## FUNDING

This work was supported by research project grants from Sapienza University of Rome (calls 2017-2020; G.L., E.C., R.N.) and by the BBSRC (BBS/E/B/000C0421; P.J.RG).

## COMPETING INTEREST

The authors have no relevant financial or non-financial interests to disclose.

## AUTHOR CONTRIBUTIONS

G.L. conceived the study and designed the experiments. A.G., P.S.N. and G.L. performed cell culture and sample preparation for endpoint analyses. A.G. and G.L performed qRT-PCR analysis. A.G. and P.S.N. performed immunofluorescence analysis. M.F. performed flow cytometry analysis. V.L. performed all the bioinformatic analysis. G.L. wrote the first manuscript draft. All authors contributed to data interpretation, critically revised the manuscript and approved its final version.

## DATA AVAILABILITY

RNA-seq data accompanying this paper are available through NCBI Gene Expression Omnibus (GEO) repository, under accession number GSE222691.

## EXPANDED VIEW FIGURE LEGENDS

**Figure EV1.**
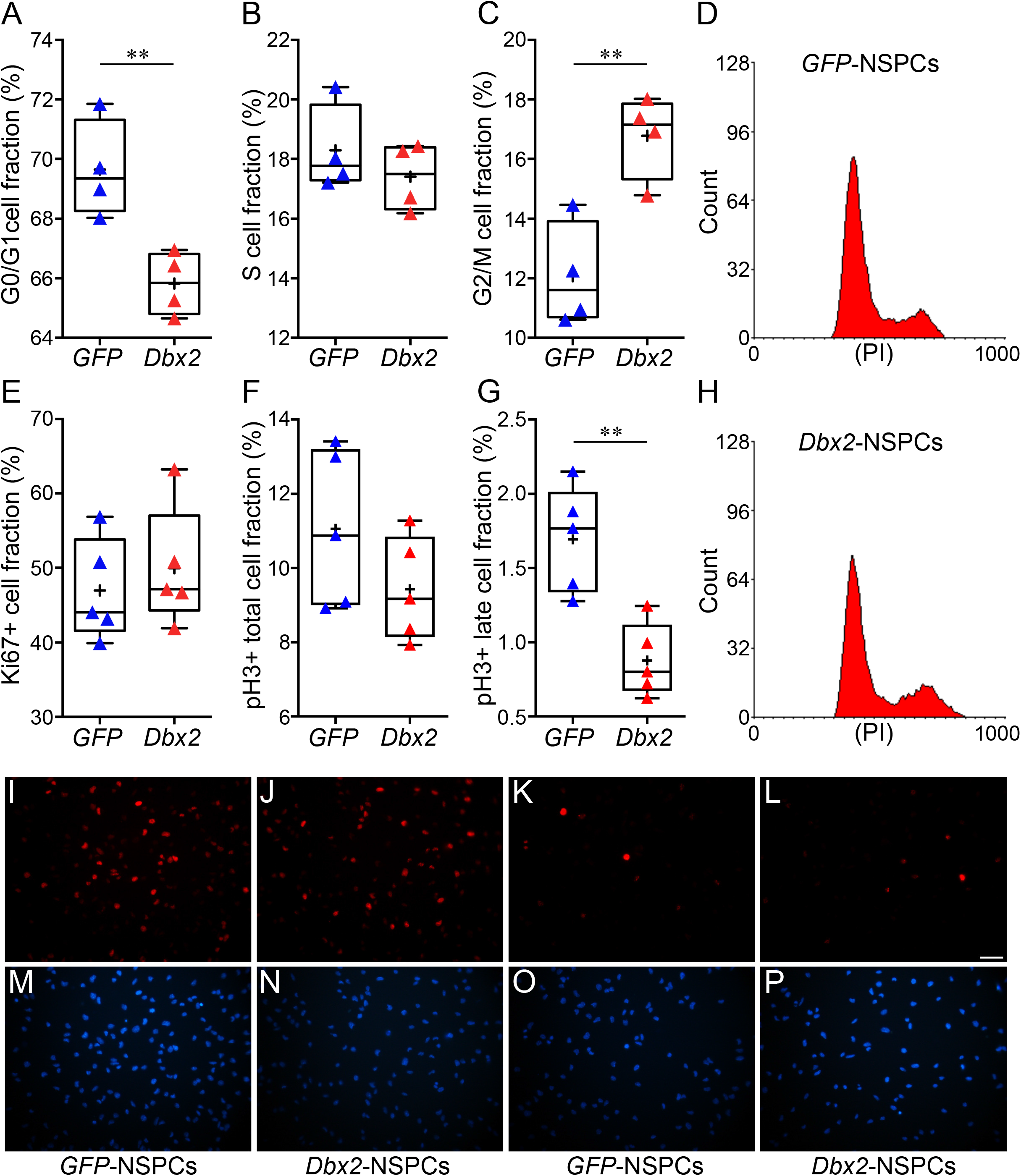
Validation of the effects of constitutive *Dbx2* overexpression in an independent pair of *GFP*-NSPCs and *Dbx2*-NSPCs. **(A** to **C)** Box-and-whisker plots of the fraction of *GFP*-NSPCs (blue triangles) and *Dbx2*-NSPCs (red triangles) in the G0/G1 **(A)**, S **(B)** and G2/M **(C)** phases of the cell cycle, after culture of an independent pair of transgenic NSPCs (n=4); **, p < 0.01, Student’s t-test. Flow cytometry histograms of PI stained *GFP*-NSPC and *Dbx2*-NSPC cultures from a representative experiment are shown in **(D)** and **(H)**, respectively. **(E** to **G)** Box-and-whisker plots of the fraction of Ki67+ **(E)**, pH3+ total **(F)** and pH3+ late **(G)** cells, after culture of an independent pair of *GFP*-NSPCs (blue triangles) and *Dbx2*-NSPCs (red triangles) (n=5); **, p < 0.01, Student’s t-test. **(I** to **P)** Representative images of *GFP*-NSPC **(I**, **M**, **K**, **O)** and *Dbx2*-NSPC **(J**, **N**, **L**, **P)** cultures stained with anti-Ki67 **(I**, **J)** or anti-pH3 **(K**, **L)** antibodies. Hoechst nuclear staining is shown in **(M** to **P)**. Scale bar, 40 μm.

**Figure EV2.**
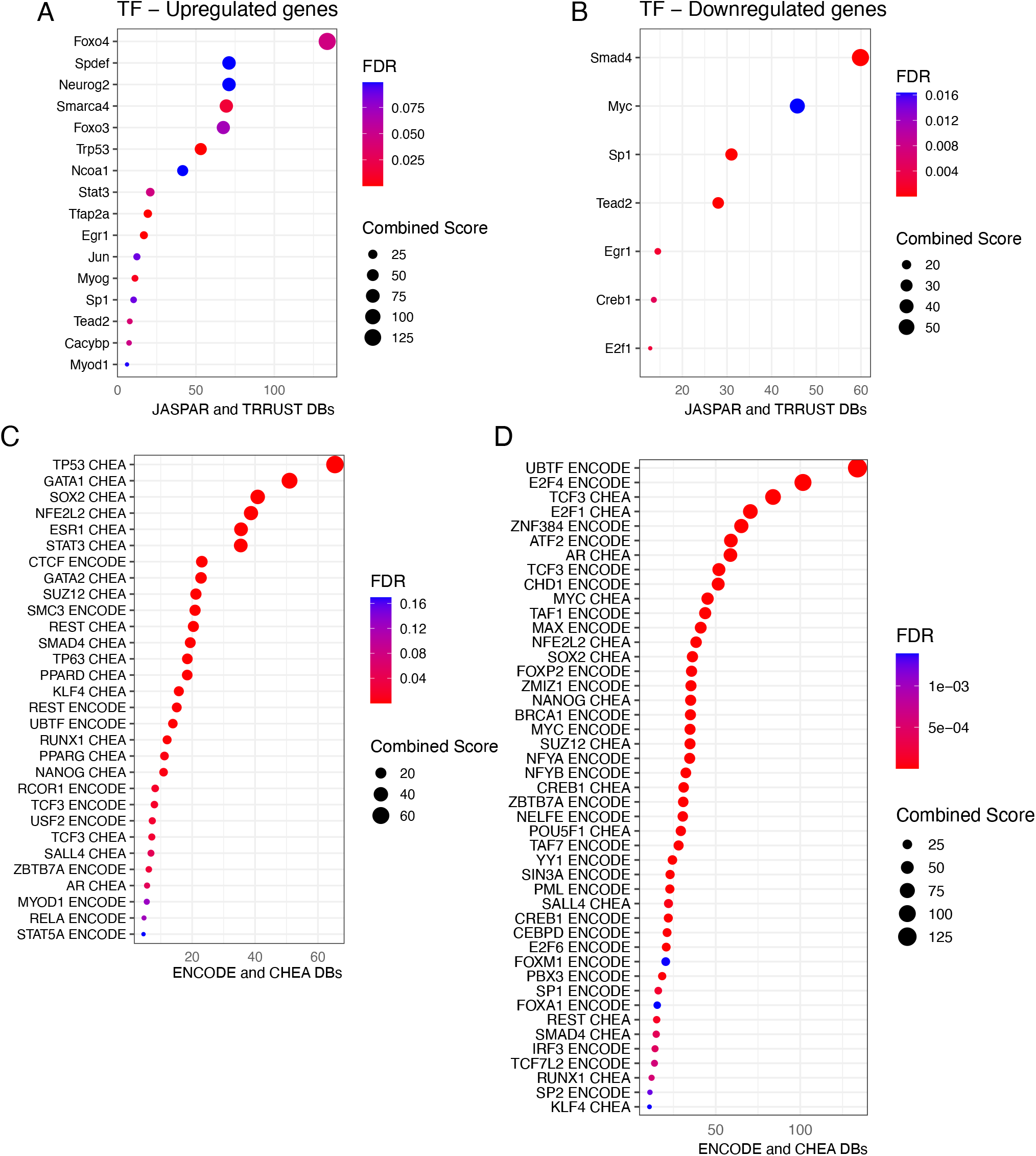
Transcription factor motifs enriched in *Dbx2*-modulated genes. (**A** to **D**) Dot plots showing the transcription factor motifs enriched in the upregulated (**A**, **C**) and the downregulated (**B**, **D**) DEGs in *Dbx2*-NSPCs vs *GFP*-NSPCs, according to the JASPAR and TRRUST databases (**A**, **B**) or the ENCODE and CHEA databases (**C**, **D**). The size of the dots is based on the count of the DEGs correlated with each motif; the colour of the dots shows the qvalue associated with gene enrichment for each motif.

**Figure EV3.**
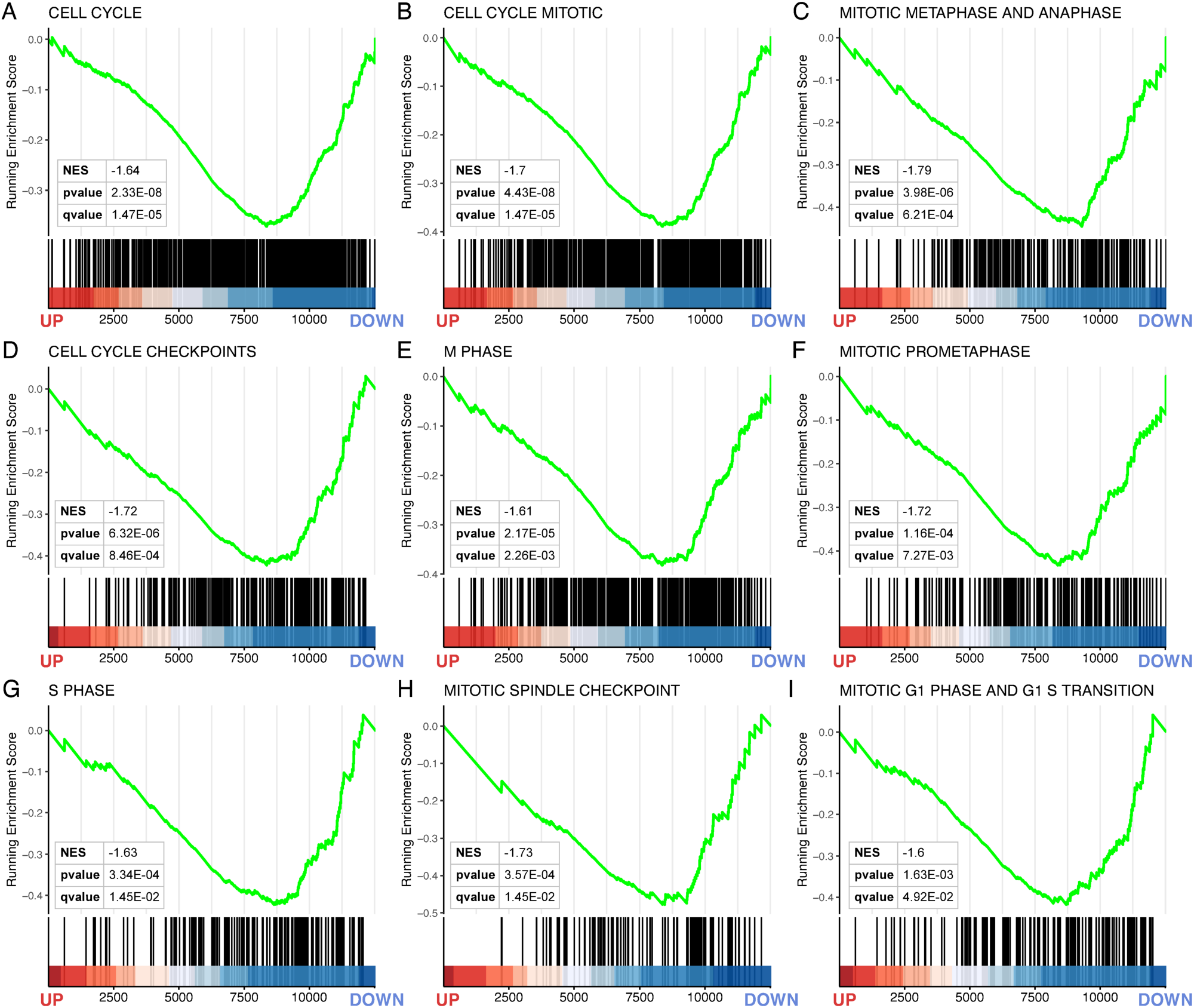
*Dbx2* overexpression inhibits the expression of gene sets associated with different cell cycle phases. (**A** to **I**) Enrichment plots showing the enrichment in the *Dbx2*-associated signature of gene sets related to different cell cycle phases, which were obtained from the “C2” curated gene sets collection in the Molecular Signatures Database, according to GSEA. Vertical black lines indicate individual members of each gene set; the heat maps at the bottom of the plots show the position of each gene within the ranked *Dbx2*-associated signature

**Figure EV4.**
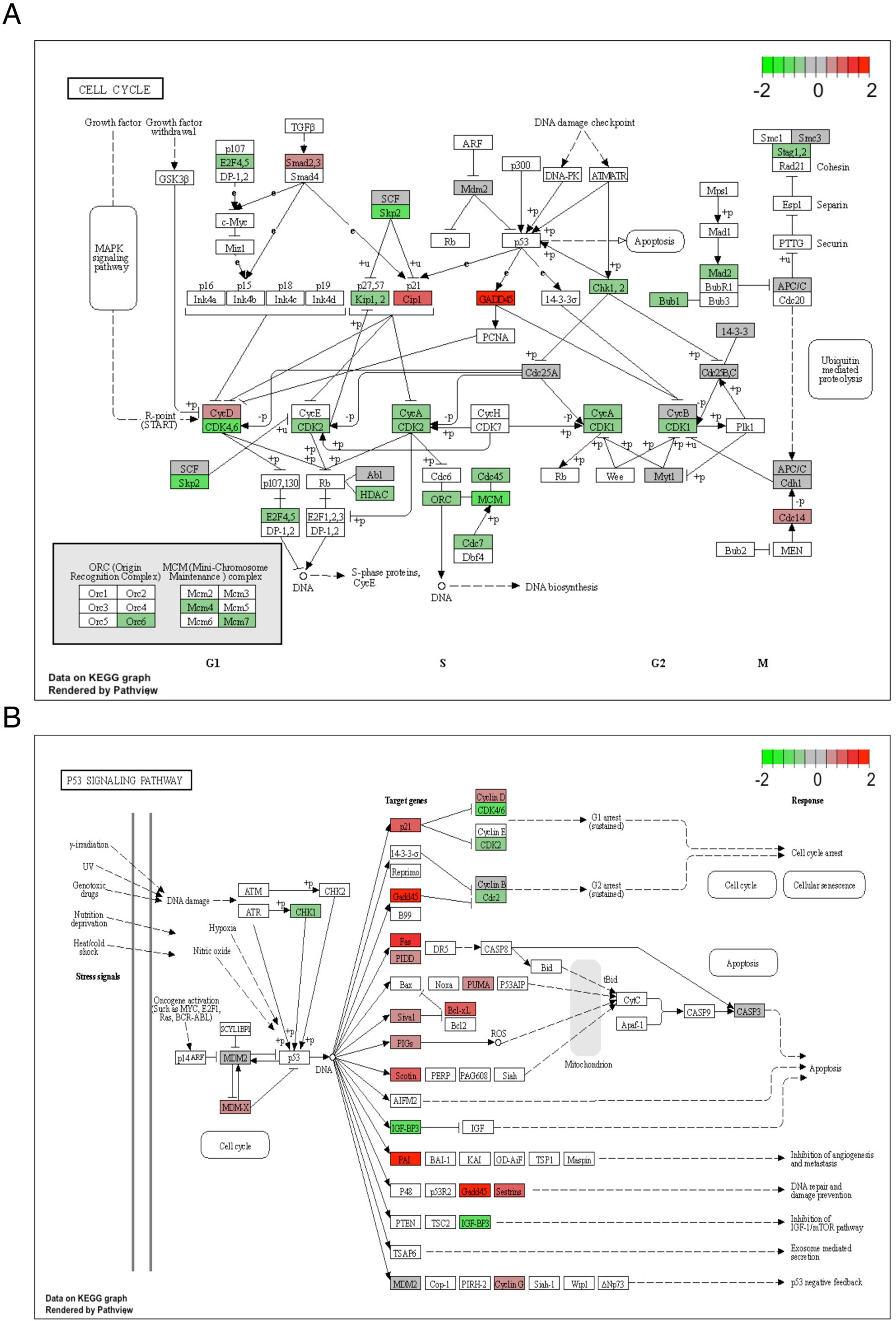
*Dbx2* overexpression causes the downregulation of gene involved in cell cycle progression and the upregulation of genes involved in cell cycle inhibition. **(A**, **B)** Pathview-generated diagrams of the cell cycle molecular pathway (**A**), or the p53 signaling pathway (**B**), according to KEGG database, showing that genes with crucial roles in cell cycle progression are downregulated in *Dbx2*-overexpressing NSPCs (green colour, FC < 0), whereas genes coding for key cell cycle inhibitors are upregulated following *Dbx2* overexpression (red colour, FC < 0).

**FIGURE EV5.**
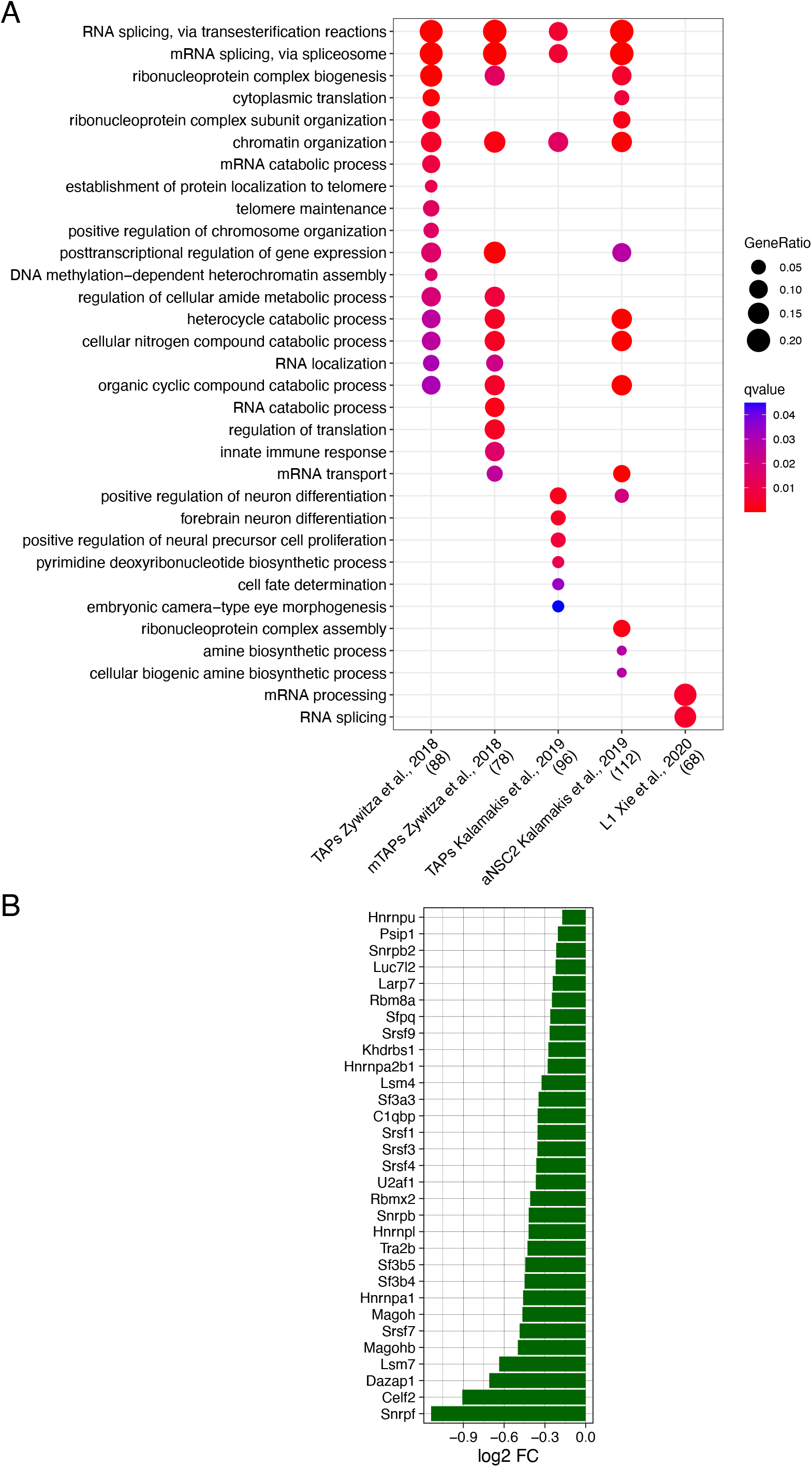
*Dbx2* affects the expression of gene sets related to both RNA processing and neurogenic lineage progression. (**A**) Dot plot showing the GO terms enriched in the gene sets obtained by subtracting cell cycle genes, taken from the KEGG and REACTOME cell cycle gene sets, from the indicated NSPC signatures [TAPs and mTAPs, (Zywitza *et al*, 2018); TAPs and aNSC2 (Kalamakis *et al*, 2019); L1 (Xie *et al*, 2020)], then selecting the downregulated DEGs in *Dbx2*-NSPCs vs *GFP*-NSPCs in the resulting gene lists. The size of the dots is based on the count of the DEGs correlated with each term; the colour of the dots shows the qvalue associated with gene enrichment for each term. (**B**) Bar plot showing the FC of genes associated with the GO terms “RNA splicing, via transesterification reactions” and “mRNA splicing, via spliceosome”, which are also associated with the NSPC signatures indicated in (**A**) and downregulated in *Dbx2*-NSPCs.

